# Icarust, a real-time simulator for Oxford Nanopore adaptive sampling

**DOI:** 10.1101/2023.05.16.540986

**Authors:** Rory Munro, Alexander Payne, Matthew Loose

**Author notes:** **Correspondence** Matthew Loose -.

## Abstract

1

**Summary:** Oxford Nanopore Technologies (ONT) sequencers enable real-time generation of sequence data, which allows for concurrent analysis during a run. Adaptive sampling leverages this real-time capability *in extremis*, rejecting or accepting reads for sequencing based on assessment of the sequence from the start of each read. This functionality is provided by ONT’s software, MinKNOW (https://community.nanoporetech.com/downloads). Designing and developing software to take advantage of adaptive sampling can be expensive due to the cost of using real sequencing consumables, using precious samples and preparing sequencing libraries. MinKNOW addresses this issue in part by providing the ability to replay previously sequenced runs for testing, using its playback functionality. However, as we have illustrated here, the sequencing output only partially changes in response to adaptive sampling instructions. Here we present Icarust, a tool enabling more accurate approximations of sequencing runs. Icarust recreates all the required endpoints of MinKNOW to perform adaptive sampling and writes output compatible with current base-callers and analysis pipelines. Icarust is capable of serving nanopore signal to simulate a MinION or PromethION flow cell experiment from any reference genome using either R9 or R10 pore signal. We show that simulating sequencing runs with Icarust provides a realistic testing and development environment for software exploiting the real-time nature of Nanopore sequencing.

**Availability and Implementation:** All code is open source and freely available here - https://github.com/LooseLab/Icarust. Icarust is implemented in Rust, with an easy to install and use docker container available.

**Supplementary Information:** Supplementary information is available here - https://github.com/Adoni5/Icarust_paper.

## 2 Introduction

Nanopore sequencing, as developed by Oxford Nanopore Technologies (ONT), is unique as read data are available for analysis as soon as individual molecules complete passing through the nanopore (Hook and Timp 2023). It is even possible to observe and analyse molecules as they are being sequenced (Loose, Malla, and Stout 2016). These properties enable methods such as adaptive sampling whereby individual molecules can be chosen for sequencing from a library (Payne, Holmes, Clarke, et al. 2021). On target molecules are left to finish passing through the pore, whereas off target molecules are rejected from the pore, in a process known as unblocking, freeing up the pore to potentially sequence more on target molecules. Real-time analysis of sequence data is possible, but tools utilising these approaches can be complex and costly to develop. Additionally, any tool changing the output of a sequencer could, unintentionally, negatively impact sequencing performance. As such, thorough testing of tools is required.

ONT provides a piece of software, MinKNOW, to control nanopore sequencing. All interactions with the sequencer are conducted via MinKNOW through an open Application Programming Interface (API) (https://github.com/nanoporetech/minknow_api). MinKNOW can be configured to playback prerecorded sequencing data, which enables some simulation options, providing a route to develop tools and analysis methods without significant expense (Munro, Santos, et al. 2022). However, these simulations are limited as reads rejected from a pore are not actually removed from the simulation - instead the original read is fragmented at the point the read would have been unblocked. This results in single original long reads being divided into smaller fragments if they are sent an unblock command. This approach also requires a pre-recorded bulk file for a given sample and experiment (Payne, Holmes, Rakyan, et al. 2018). Other simulation tools have been developed to address similar problems. Uncalled, software to implement adaptive sampling functionality, included a simulator that can be driven from pre-existing sequence fast5 files (Kovaka et al. 2021). However, to our knowledge this simulator does not remove the need for pre-existing data sets nor can any generic implementation of adaptive sampling be developed to use it. Other ONT signal simulators exist including Squigulator and DeepSignal 1.5 (Gamaarachchi et al. 2023; Li et al. 2020), although these approaches do not enable real-time simulation for serving squiggle chunks for adaptive sampling.

To address these challenges, we developed Icarust, a simulator for nanopore sequencing, which mimics the action of MinKNOW. Icarust generates signal data (“squiggle”) derived from the output of “Scrappie” (https://github.com/nanoporetech/scrappie) and serves this signal in real-time using an identical API to the one implemented in MinKNOW itself. Icarust can also generate R10.4 pore data using models provided by ONT (https://github.com/nanoporetech/kmer_models/). Icarust can simulate any genome, incorporate barcodes and respond to requests to reject reads via the same adaptive sampling API as MinKNOW. Icarust can also simulate amplicon based sequencing experiments. Data from Icarust are written to files (currently FAST5 only) which are compatible with Oxford Nanopore base-callers. Importantly, this tool is not intended to be used to explore base-calling itself. None of the signal methods employed will capture any of the nuance of real signal, rather they are simply signals that can be meaningfully interpreted by base-callers. This tool enables cheap testing and development of software designed to exploit adaptive sampling and real-time analysis of nanopore sequence data.

## 3 Software Implementation

Icarust is implemented in Rust (https://www.rust-lang.org/), using the tonic package (https://github.com/hyperium/tonic/tree/master) to provide gRPC Remote Procedural Call (gRPC) support. Rust was chosen as it was the fastest gRPC implementation (https://github.com/LesnyRumcajs/grpc_bench/wiki/2022-01-11-bench-results), and is naturally asynchronous. All MinKNOW API endpoints required by ReadFish (Payne, Holmes, Clarke, et al. 2021) are implemented.

### 3.1 Squiggle generation

In order to serve signal data (squiggle), we employ one of two methods. The first method pre-computes all squiggle from a user provided reference genome as the conversion from sequence to squiggle is too slow to be performed in real-time. For R9 data, we utilise the ONT base-caller Scrappie to convert the reference to an array of squiggle values. Icarust then randomly selects a read length and starting location from the pre-computed array for each channel, and serves chunks of this selection, at approximately 4kHz. This approach is useful as the signal is generated by reversing the base-caller network, but is limited by the large file size of the pre-computed squiggle and the lack of support for later pore types in Scrappie.

The second method utilises pore models provided by ONT for research purposes. We again select a random start point and read length from the reference sequence and then convert this sub sequence into signal by sliding an appropriate base window along the sequence and converting the data into signal measurement samples. The values listed in the pore models are z-score normalised, as such some manipulation is needed to de-normalise the values back into a form that can be called by the base-caller. This de-normalisation is shown in Equation (1)

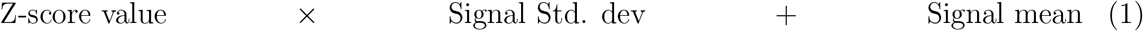

These values can be changed, however the values chosen here are sufficient to get signal that can be base-called accurately enough for alignment. This way it is possible to directly convert sequence at the start of the simulation, so no pre-computation is required. This signal is far from perfect, however our aim is to simply test the real-time feedback properties of sequencing, not simulate signal perfectly.

### 3.2 Barcoding

In order to simulate multiplexed samples, the sequences for the NB12 barcoding kit (ONT) were converted to squiggle. The correct complements are then added to the start and end of the squiggle for a read, and are padded with some “adaptor” signal. The ratio of barcodes in a sample can be altered by providing weights to each barcode in a simulation profile TOML.

### 3.3 Flow cell health modelling

To more accurately recreate the decline of flow cell health across a sequencing run and capture the effects of applying adaptive sampling, we assign each simulated “channel” a probability to become saturated and so unavailable for further sequencing. When simulation begins, some channels are labelled as saturated immediately (default 15%). After a read ends, either naturally or as a result of an unblock, we determine if the channel can produce another read. The probability of this is determined by the base chance (described below) multiplied by the length of the read, divided by 10 000. This takes into account the fact that longer reads are more likely to block a channel than shorter reads. The base chance for saturation is a function of the number of available channels, the desired total yield and the mean read length as seen in eq. (2).

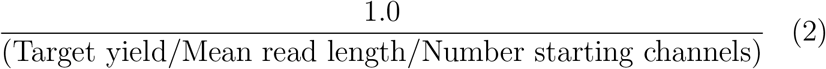

These parameters provide a near realistic decay in sequencing performance that mimics the overall total yield of a genuine sequencing experiment. All parameters can be defined by the user (see below).

### 3.4 Configuration

Icarust is configured by a combination of two files. A configuration INI file provides Icarust software specific configuration, such as where TLS certificates can be found for securing the gRPC connection between the MinKNOW API and Icarust, and what ports Icarust listens on. It also provides sequencer specific configuration, such as the number of channels being simulated.

A second configuration TOML file holds the settings about the specific simulation taking place. This “Simulation profile” contains tables that can alter variables about the run, such as the proportions of the species being sequenced, the average read length of the sample, which barcodes are present on each sample. A full list of settings is detailed in the source code README (https://github.com/Adoni5/Icarust/blob/master/README.md).

### 3.5 Output

Akin to a typical nanopore sequencing run, when 4 000 reads have finished sequencing, a FAST5 file containing the read squiggle and metadata is written out. The directory structure for the FAST5 files mimics the same structure employed by MinKNOW. The availability of these files allows for downstream post run analysis of the simulation.

## 4 Results and Discussion

### 4.1 Example use case

In order to test the utility of our tool, we compared running adaptive sampling using play-back and Icarust with the R9.4 pore Scrappie model. We performed the simple experiment recommended by ReadFish, targeting Chromosome 20 and 21 on the human genome. This resulted in four runs in total. A control for playback and Icarust and then an adaptive sampling run using playback and Icarust. As shown in Figure 1a, the application of adaptive sampling worked for each run simulation method, with the median read lengths reduced for each off target Chromosome when ReadFish is applied. The median read lengths for the on target chromosomes (Chromosomes 20 and 21) remained close to or the same as the control for both simulation methods.

**Figure 1:**
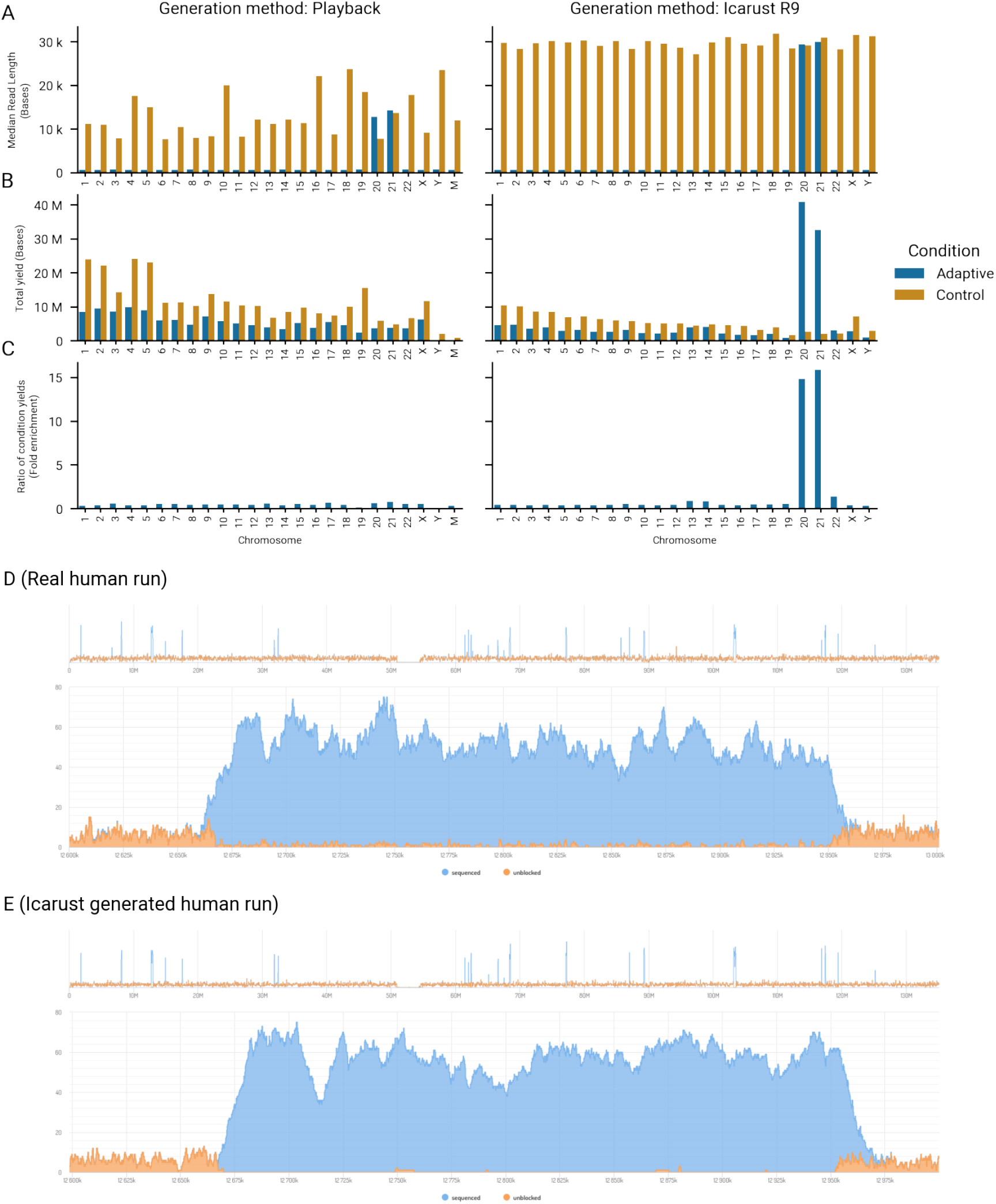
Comparison of an adaptive sampling experiment, targeting Chromosomes 20 and 21 on the human genome, between two simulated runs. The left column visualises data generated using MinKNOWs playback, the right column data generated using Icarust with R9 data. The two run conditions, Control (orange) or Adaptive sampling (blue) are shown. A). The median read length of the two runs for each condition. B). The total yield in bases of the two runs for each condition. C). The difference between the yield in the Adaptive and Control conditions as the fold change for each run. D,E). Coverage plots showing the same adaptive sampling target (Chr 11, 12,666,921 - 12,952,237) from a real human sequencing run, and a replication run, simulated using Icarust R9 data. The top section of the plots shows the whole coverage over Chr 11. Coverage is split between Sequenced (Blue) and Unblocked reads (Orange). Charts plotted using MinoTour.

However, the effect adaptive sampling has on the experimental outcome is varied, shown in Figures 1b and 1c. The yield (Figure 1b) for playback runs is lowered when adaptive sampling is applied. This is likely a consequence of the unblocking of reads without the ability to replace them with a new molecule. As a consequence, there is no enrichment in yield. However, in the Icarust generated experiment, where reads can be replaced with new molecules, we see clear enrichment of the target chromosome. This is more clearly shown in Figure 1c, where we can see the ratio of the yields between the Control and Adaptive conditions. In the run simulated by Icarust, we see the on target yield enriched by approximately 15 ×, whereas in the MinKNOW simulated run we see a fall in yield to about 0.7 × of the control yield.

In a separate simulation, we aimed to recreate a previously performed human adaptive sampling sequencing run using Icarust. We ran Icarust using the same target set as the original run, and used ReadFish to perform adaptive sampling. We then uploaded the run to MinoTour in real-time. We found that we were able to recreate a reasonable approximation of the run, as shown in Figures 1d and 1e.

## 5 Conclusion

Here we present Icarust, a tool designed to allow easy testing of Adaptive Sampling software and experimental setups for ONT sequencers. Icarust allows users to quickly and cheaply test adaptive sampling experiments and develop new software for ONT adaptive sampling workflows. Icarust can simulate barcoded and non barcoded sequencing runs from any provided reference sequence, sequencing runs using amplicon based libraries (Munro, Holmes, et al. 2023), and can simulate both MinION or PromethION scale flow cells with either R9 or R10 pore signal.

Implemented in Rust, Icarust is fast, reliable, memory safe and energy efficient. Whilst Icarust is currently limited to only writing FAST5 files, work is ongoing to implement POD5 support. Icarust is not intended to be a perfect recreation of real squiggle production, instead it is a close enough facsimile to allow software development and testing of experiment setup. The software is freely available, with a maintained docker image allowing easy adoption by the Nanopore Community.

## 6 Acknowledgements

We would like to thank all of the Deepseq Nottingham and Nadine Holmes for library preparation, and Lukas Weilguny and Lea Kaufmann for helpful troubleshooting and discussions.

## Notes

https://github.com/Looselab/Icarust

### Competing Interest Statement

Matthew Loose was a member of the MinION access program and has received free flow cells and sequencing reagents in the past.
ML has received reimbursement for travel, accommodation and conference fees to speak at events organised by Oxford Nanopore Technologies.

